# Anaplerotic pathways in *Halomonas elongata*: the role of the sodium gradient

**DOI:** 10.1101/2020.05.13.093179

**Authors:** Karina Hobmeier, Marie C. Goëss, Christiana Sehr, Hans Jörg Kunte, Andreas Kremling, Katharina Pflüger-Grau, Alberto Marin-Sanguino

## Abstract

Salt tolerance in the *γ*-proteobacterium *Halomonas elongata* is linked to its ability to produce the compatible solute ectoine. The metabolism of ectoine production is of great interest since it can shed light on the biochemical basis of halotolerance as well as pave the way for the improvement of the biotechnological production of such compatible solute. The ectoine production pathway uses oxaloacetate as a precursor, thereby connecting ectoine production to the anaplerotic reactions that refill carbon into the TCA cycle. This places a high demand on these reactions and creates the need to regulate them not only in response to growth but also in response to extracellular salt concentration. In this work we combine modeling and experiments to analyze how these different needs shape the anaplerotic reactions in *H. elongata*. First, the stoichiometric and thermodynamic factors that condition the flux distributions are analyzed, then the optimal patterns of operation for oxaloacetate production are calculated. Finally, the phenotype of two deletion mutants lacking potentially relevant anaplerotic enzymes: Phosphoenolpyruvate carboxylase (Ppc) and Oxaloacetate decarboxylase (Oad) is experimentally characterized. The results show that the anaplerotic reactions in *H. elongata* are indeed subject to different evolutionary pressures than those of other gram-negative bacteria. Ectoine producing halophiles must meet a higher metabolic demand for oxaloacetate and the reliance of many marine bacteria on the Entner-Doudoroff pathway compromises the anaplerotic efficiency of Ppc, which is usually one of the main enzymes fulfilling this role. The anaplerotic flux in *H. elongata* is contributed not only by Ppc but also by Oad, an enzyme that has not yet been shown to play this role *in vivo*. Ppc is necessary for *H. elongata* to grow normally at low salt concentrations but it is not required to achieve near maximal growth rates as long as there is a steep sodium gradient. On the other hand, the lack of Oad presents serious difficulties to grow at high salt concentrations. This points to a shared role of these two enzymes in guaranteeing the supply of OAA for biosynthetic reactions.

## 1 Introduction

The halophilic *γ*-proteobacterium *Halomonas elongata* DSM 2581^*T*^ has a broad salt tolerance and can even grow in salt saturated brines (*>* 30% NaCl) [35] thanks to the accumulation of the compatible solute ectoine, which can reach molar concentrations in the cytoplasm without disrupting cellular processes. The synthesis of ectoine from its precursor oxaloacetate (OAA) withdraws a high amount of carbon out of the TCA cycle, which must then be replenished by anaplerotic reactions. This makes the PEP-Pyr-OAA node, already acknowledged as a major switching point for carbon metabolism [29], even more important for this organism.

Figure 1 shows the PEP-Pyr-OAA node and the glyoxylate shunt in *H. elongata* as it has been determined by genomic analysis [30, 26]. The glyoxylate shunt bypasses the two decarboxylation steps of the TCA cycle and thus enables the replenishment of C4 carbon skeletons from Acetyl coenzyme-A. This pathway allows an efficient growth on acetate but its operation on glucose as a carbon source would result in a loss of one third of the total carbon in the decarboxylation of pyruvate to Acetyl-CoA by Pyruvate DH. Growth on hexoses can be carried out efficiently thanks to the anaplerotic reactions.

**Figure 1:**
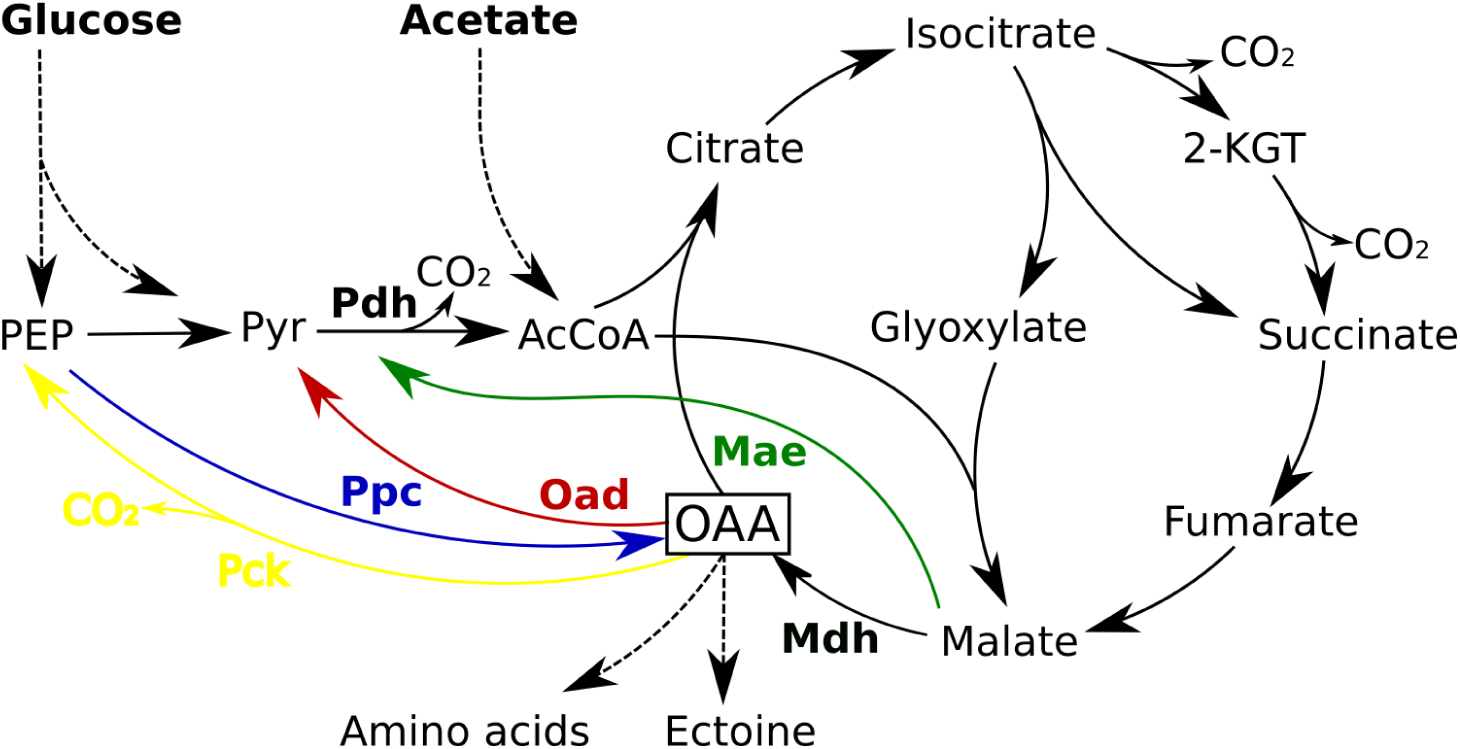
Anaplerotic reactions and TCA cycle in *Halomonas elongata*. Shown are the four enzymes PEP carboxylase (PPC, blue), Oxaloacetate decarboxylase (OAD, red), Malic enzyme (MAE, green) and Phosphoenolpyruvate carboxykinase (PCK, yellow) that have the potential to catalyze anaplerotic reactions

There are four enzymes that fullfil the stoichiometric requirements to catalyze anaplerotic reactions: PEP carboxylase (Ppc), Malic enzyme (Mae), PEP carboxykinase (Pck) and the membrane bound Oxaloacetate decarboxylase (Oad). The first of these enzymes, Ppc, has been known for a long time to carry most of the anaplerotic flux in *Escherichia coli* growing on glucose, glycerol or pyruvate [3, 24] and its regulation has been shown to play a critical role in carbon catabolism [37]. The second enzyme in the list, the malic enzyme (Mae), is present in two variants with different cofactor specificities (NADH and NADPH). The third candidate, phosphoenolpyruvate carboxykinase (Pck), is also an important enzyme in this node and catalyzes a key step for gluconeogenesis converting of OAA into PEP. Finally, Oad deserves special attention in *H. elongata*. This enzyme catalyzes the interconversion of OAA into pyruvate. Cytoplasmic versions of this enzyme are considered to be irreversible, thermodynamically unable to function in the anaplerotic sense and usually involved in gluconeogenesis [25]. In *H. elongata*, however, Oad is a membrane bound sodium pump [9] and it has been hypothesized [17] that it can operate as an anaplerotic reaction, thus using the considerable sodium gradient available under high salt concentrations to drive the carboxylation of pyruvate in *H. elongata*. To the extent of our knowledge, such a function has never been observed *in vivo*. The different physiological roles described so far for Oad go in the opposite direction, for example, as a sodium-transport system energized by the decarboxylation of OAA into pyruvate [8, 10, 6, 7] and as part of a catabolic pathway in a mutant strain of *Lactococcus lactis* [28].

The aim of this work is to shed light on the way anaplerotic reactions are organized in *H. elongata* through a combination of theoretical and experimental methods. Due to their mechanisms, the action of two of these enzymes (Oad and Ppc) cannot be distinguished through labeling experiments [23]. Therefore, we constructed mutant strains lacking each of the enzymes (Ppc and Oad) and characterized their phenotypes in comparison to the wild type. The results of this experiments are examined under the light of thermodynamic analysis and simulations of constraint based models that are customized to deal with the challenge of estimating physico-chemical magnitudes at high salt concentrations.

## 2 Material and Methods

### Strains, plasmids, and growth conditions

*H. elongata* DSM 2581^*T*^ and the mutant strains were routinely grown in minimal medium MM63 or LB medium containing 1 M NaCl at 30 ° C under shaking at 220 rpm. All *Escherichia coli* strains were grown in LB medium at 37 ° C amended with the antibiotic necessary for maintenance of the plasmid. All plasmids and genetic manipulations were performed in *E. coli* DH5*α* or *E. coli* DH5*α λ*-pir.

### Construction of plasmids pSEVA_Δ*ppc*::Sm and pSEVA_Δ*oad* ::Sm

Replacement of the *ppc* gene with *aadA*, encoding a Streptomycin resistance cassette was performed by adapting the method previously reported for *Pseudomonas putida* [20]. The plasmids were assembled by Gibson Assembly (NEB) according to the suppliers manual. Oligonucleotides used for the gene amplification are shown in table 1. A DNA fragment containing about 500 bp of the genomic sequence upstream of the target gene (*ppc* or *oad*) followed by the sequence of the *aadA* gene, encoding the streptomycin resistance cassette, and about 500 bp of the chromosomal region downstream of the target gene, was inserted into pSEVA212S, carrying the R6K suicide origin of replication [19], which had been previously linearized with EcoRI. The plasmid was transformed [5] into *E. coli* DH5*α λ*-pir and clones carrying it were selected on LB agar plates supplemented with 200 *µ*g/ml streptomycin (Sm 200). The correct assembly of pSEVA_Δ*ppc*::Sm and pSEVA_Δ*oad* ::Sm was verified by isolation of the plasmids and sequencing.

**Table 1:**
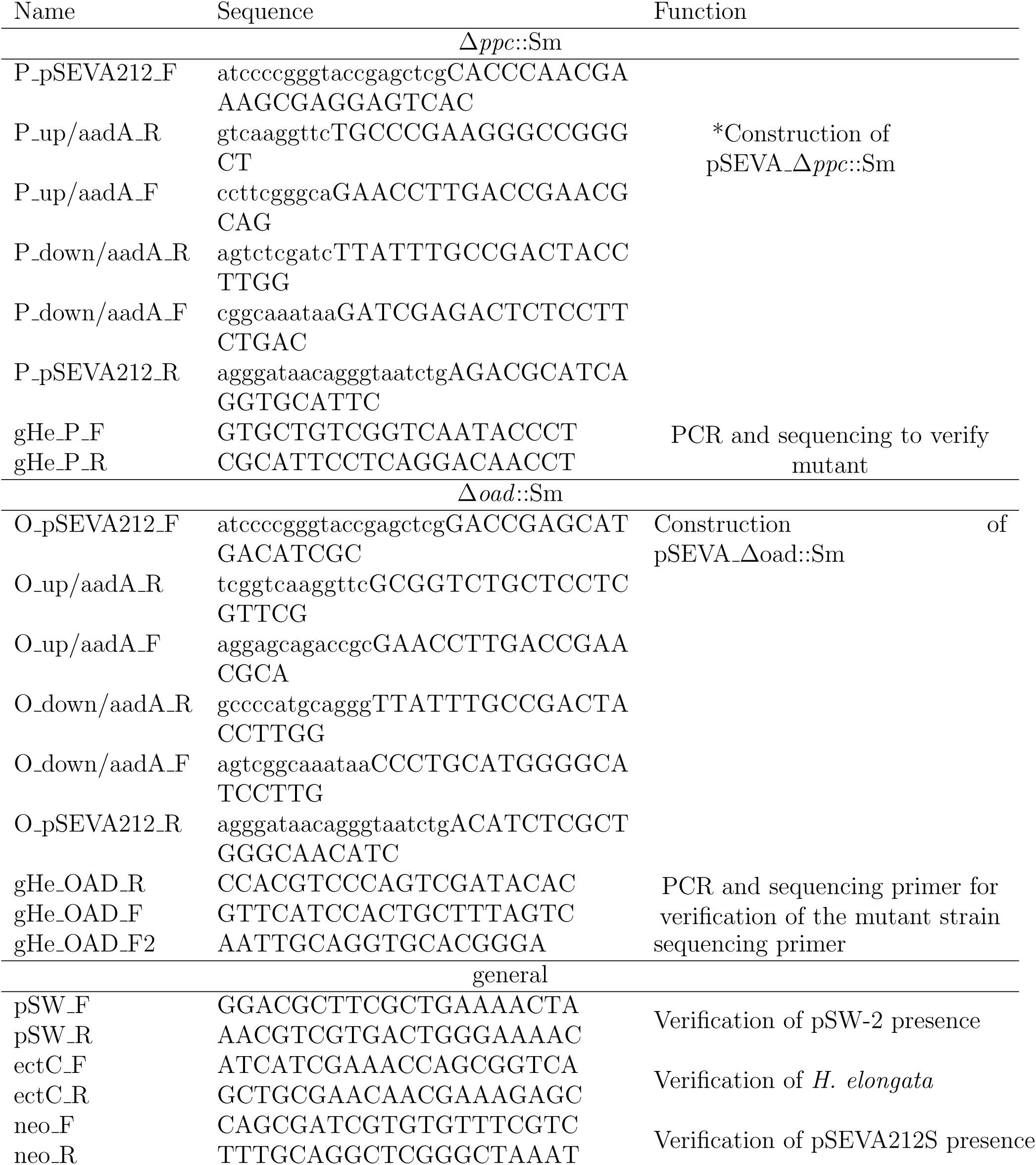
Oligonucleotides used in this work. * Sequences complementary to one fragment are shown in lower case and to the other in upper case.

### Conjugal transfer of plasmids into *H. elongata* by triparental mating

Plasmids were transferred to *H. elongata* by a two step mating procedure. First, plasmid pSW-2 encoding the 1-SceI endonuclease [20] was transferred to *H. elongata*. Therefore, the donor strain *E. coli* DH5*α λ*-pir (pSW-2), the recipient strain *H. elongata*, and the helper strain *E. coli* HB101 (pRK600) were grown in LB medium, amended with the antibiotic necessary to assure plasmid maintenance (pSW-2: 10 *µ*g/ml gentamycin and pRK600: 35 *µ*g/ml chloramphenicol) until the stationary phase (over night). Optical densities at 600 nm were measured and 1 ml of the donor culture was mixed with the amount of recipient and helper culture to obtain OD ratios of 1:1:1 and 1:2:2 respectively. These suspensions were centrifuged, the supernatant was discarded by decanting and the cell pellet was resuspended in the remaining medium rest. This condensed cell suspension was pipetted on one spot of a LB agar plate containing 0.5 M NaCl, and incubated for 5 h at 30 *°* C. Afterwards, the cells were resuspended in 600 *µ*l of a 1 M NaCl solution, an aliquot of 100 *µ*l was plated on LB agar plates containing 0.5 M NaCl, 500 *µ*g/ml ampicillin and 50 *µ*g/ml gentamycin and the plates were incubated for 48 h at 30 *°* C. Colonies were picked and the presence of pSW-2 was verified by PCR with pSW-F and pSW-R [20] (table 1).

Next, pSEVA_Δ*ppc*::Sm and pSEVA_Δ*oad* ::Sm were transferred and integrated in the genome of *H. elongata* (pSW-2) again by triparental mating. This was performed as described above but using *H. elongata* (pSW-2) as recipient, *E. coli* DH5*α λ*-pir (pSEVA_Δ*ppc*::Sm or pSEVA_Δ*oad* ::Sm) as donor and *E. coli* HB101 (pRK600) as helper. The cointegrates were selected by plating different dilutions of the mating mixture on LB agar plates containing 0.5 M NaCl, 500 *µ*g/ml ampicillin, 60 *µ*g/ml kanamycin, and 50 *µ*g/ml gentamycin to obtain single colonies.

### Resolution of cointegrates to obtain marker substitution mutants

To resolve the cointegrates a single colony was picked from the selective agar plate and grown in 3 ml LB medium with 1 M NaCl, 500 *µ*g/ml ampicillin, 200 *µ*g/ml streptomycin and 50 *µ*g/ml gentamycin over night at 30 *°* C. The next day, cells were pelleted by centrifugation, washed once in LB medium containing 1 M NaCl, and subsequently resuspended to an OD600 of 1 with LB medium with 1 M NaCl, 500 *µ*g/ml ampicillin, and 50 *µ*g/ml gentamycin. To four ml of that culture, 80 *µ*l of 3-methylbenzoate were added to a final concentration of 10 mM to induce the expression of 1-SceI and cells were incubated at 30 *°* C under shaking.

After 3.5 h, an aliquot of cell suspension was plated on LB agar with 1 M NaCl, 500 *µ*g/ml ampicillin and 200 *µ*g/ml streptomycin and incubated at 30 *°* C for 48 hours. Single colonies were picked and gridded on LB plates with 1 M NaCl, 500 *µ*g/ml ampicillin, and either 60 *µ*g/ml kanamycin or 200 *µ*g/ml of streptomycin to identify those clones, that lost the kanamycin resistance cassette but maintained the streptomycin cassette. Clones showing the desired phenotype should represent the desired mutants. This was verified by PCR using primers (gHe_P_F, gHe_P_R, gHe_OAD_F, gHe_OAD_R, gHe_OAD_F2 ; table 1) that hybridized to sequences outside the 500 bp flanks used in pSEVA_Δ*ppc*::Sm or pSEVA_Δ*oad* ::Sm and sequencing of the whole fragment.

The two mutant strains of *H. elongata* obtained by this procedure: Δ*ppc*::Sm and Δ*oad* ::Sm will be named *H. elongata*-PPC and *H. elongata*-OAD respectively.

### Growth experiments on glucose or acetate in microplates

The growth experiments were performed in MM63 medium with 27.75 mM glucose or 27 mM acetate, respectively. Initially, a single colony of each strain was picked per replicate and grown in 3 ml LB medium with 1 M NaCl. The next day, an aliquot was used to inoculate 3 ml of MM63 medium with 1 M NaCl. From this culture, three aliquots were used to inoculate each of the pre-cultures for the growth experiment in MM63 medium with either 0.17 M, 0.5 M, 1 M, or 2 M NaCl. Inoculation was always done in a way to reach an initial OD600 of 0.01 and incubation was in all cases at 30 *°* C in the shaker until the late exponential or early stationary phase. For the growth experiment, 200 *µ*l of medium (MM63 with 0.17 M, 0.5 M, 1 M, or 2 M NaCl) were inoculated with 2 *µ*l of the respective pre-culture adapted to the respective NaCl concentration in a sterile 96-well microtiter plate with optical bottom (Greiner, Germany) and incubated in an automated microplate reader (Tecan, Austria) at 30 *°* C. Optical density was measured every 10 min after vigorous shaking for 3 sec over a period of 16 to 22 h. In a first round of experiments, each condition was measured at least 3 times, i.e. starting with 3 different colonies per strain and per carbon source. The maximum growth rate of each well was defined as the maximum value of linear regression of the logarithm of the OD along a sliding window of at least 15 points. In order to ensure that the chosen values were part of a sustained exponential growth phase, only values that had a determination coefficient above 0.9 were considered eligible. Conditions that yielded significant differences in the first round of screening were analyzed in a second round using four biological replicates (from different colonies) each having two technical replicates (two wells inoculated from the same preculture). Differences reproduced in both experiments were tested in flasks.

### Growth experiments on glucose or acetate in shake flasks

Growth experiments in shake flasks were performed in MM63 medium with 27.75 mM glucose or 27 mM acetate, as carbon source in the presence of 0.17 M NaCl. Again, a single colony of the relevant strain was picked per replicate and grown in LB medium with 1 M NaCl. From these cultures aliquots were taken to inoculate the pre-cultures in MM63 medium with 0.17 M NaCl to an OD600 of 0.01. Pre-cultures of *H. elongata* wt were incubated for about 12 h until they reached an OD600 of 1.5 to 2.6 on glucose and 0.5 to 0.8 on acetate, whereas pre-cultures of *H. elongata*-PPC were incubated for approximately 48 h to reach an OD between 4.9 and 6.0 in case of growth on glucose and between 1.0 and 1.2 in case of acetate grown cells. From theses pre-cultures pre-warmed 25 ml of MM63 medium with 0.17 M NaCl and either glucose or acetate as carbon source were inoculated to an OD600 of 0.05 and incubated under shaking for 11 h. The OD600 was measured every hour in a spectrophotometer (Eppendorf, Deutschland). The experiment was performed in triplicates, i.e. starting from 3 different colonies per strain.

### Growth experiments on glucose or acetate in shaking flasks for ectoine quantification

The cultivations were performed in MM63 minimal medium with 27.75 mM glucose or acetate as carbon source in the presence of 2 M NaCl. For each strain (wildtype, *H. elongata*-PPC and *H. elongata*-OAD) three single colonies were picked and grown in 1 M NaCl LB medium. From these precultures aliquots were taken to inoculate accommodation cultures to an OD600 of 0.01 in MM63 medium with 1 M NaCl and glucose (or acetate) as sole carbon source. This preculture step was then repeated once more with 2 M NaCl MM63 minimal medium. From these precultures 50 mL of MM63 medium with 2 M NaCl and either glucose or acetate were inoculated to an OD600 of 0.01 and incubated under shaking 220 rpm for up to 70 hours. The OD600 was measured regularly in a spectrophotometer (Eppendorf, Deutschland) until cells were harvested during the exponential phase. To release the intracellular ectoine a two-phase extraction protocol developed by Bligh and Dyer [4] and further modified by Galinski and Herzog [13] was applied. For this, the samples were centrifuged (15000 g, 25 *°* C) for 5 min and the supernatant was removed. The remaining cell pellet was resuspended in 250 *µ*L extraction solution (methanol, chloroform, bidest. water in ratios 10:5:4). After an incubation period of 30 min under shaking (300 rpm, Thermomixer Comfort Typ 5, Eppendorf) at room temperature 65 *µ*L chloroform and bidest. water were added and incubated a second time under shaking (300 rpm, Thermomixer Comfort Typ 5, Eppendorf) at room temperature for 10 min. To achieve the desired phase separation the samples were centrifuged once more at 8000 *x g* for 5 min at room temperature. Then, 100 *µ*L from the 250 *µ*L aqueous upper phase were taken and diluted 1:10 using the mobile phase (Acetonitrile/phosphate buffer). Ectoine was measured by absorbace at 210nm after elution through a Nucleodur 100-5 NH2-RP CC 125/4 column (Macherey & Nagel).

### Thermodynamics-based Metabolic Flux Analysis (TMFA)

A simple model of the TCA cycle and the anaplerotic pathways was formulated based on previous stoichiometric models [17] and additional thermodynamic information [2, 12]. The stoichiometric and thermodynamic constraints were combined within the well establish TMFA framework [15, 16] into a Mixed Integer Linear Program (MILP) that enables the calculation of optimal flux distributions. Two modifications were introduced to the standard TMFA methodology. First, an additional constraint was added to ensure that the thermodynamic driving forces of all reactions were higher in magnitude than a pre-established limit. This becomes the Minimum Driving Force (MDF) as defined elsewhere [22]. Second, a necessary step for thermodynamic calculations is the calculation of metabolite activities. The activity of a metabolite is a function of its concentration *a*_*i*_ = *γ*_*i*_ *c*_*i*_ where *γ*_*i*_ is the activity coefficient. Activity coefficients in biochemistry are often calculated using the Debbye-Huckel equation [2], at *T* = 25*°C*. 

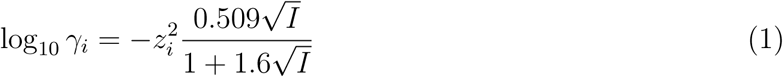

At salt concentrations above 0.3 Molar, the D-H equations are no longer able to provide valid estimates for the activity coefficient because short range ion-specific interactions become significant, the effect of specific ion can be accounted for using methods such as Specific Interaction Theory (SIT) [14]. This model requires the use of interaction coefficients for all the relevant ions *k* = 1 *… N*_*k*_: 

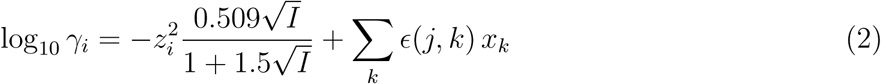

At *I >* 0.25*M*, we use equation [14]

cations 

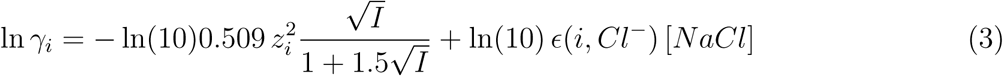

anions 

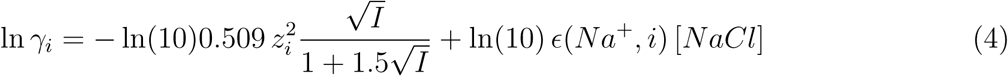

protons 

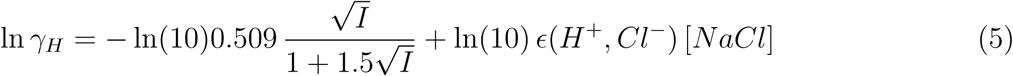

The most relevant coefficients in this case are *ϵ*(*Na*^+^, *Cl*^−^) = 0.03 and *ϵ*(*H*^+^, *Cl*^−^) = 0.12.

So every calculation involving extracellular species has to be performed according to SIT. The usual formula to transform energies of formation [2], for instance, has to be corrected:

cations 

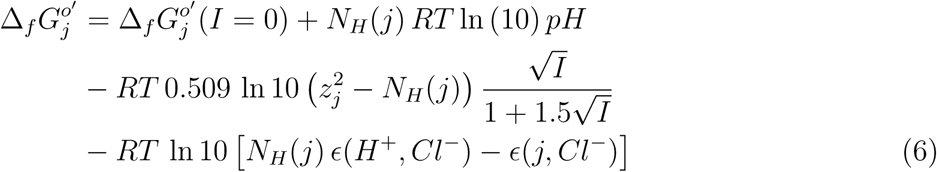

anions 

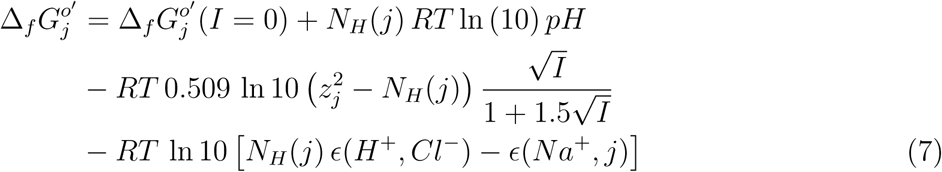

Once the model was formulated, TMFA simulations were performed at salt concentrations 0.17 M, 0.5 M, 1 M and 2 M. For each case, ATP or oxaloacetate production were maximized in order to explore the potential theoretical yields. The simulations were repeated imposing the additional restriction of the minimal driving force (MDF) being higher than different thresholds. The thresholds for MDF were chosen to ensure enzyme efficiencies ranging from 1 % to 99 % at the thermodynamic bottleneck. For simulations maximizing the yield of oxaloacetate, a minimum ATP production of 4.5 mmol / g DW was imposed to obtain a realistic split of the metabolic flux between oxaloacetate synthesis and catabolic TCA cycle of roughly 1:2 [27]. This choice does not affect the absolute differences in OAA yield between different solutions, but filters out wasteful solutions in terms of ATP.

## 3 Results

### 3.1 Ectoine accumulation dramatically increases the demand for Oxaloacetate

The concentration of ectoine in the cytoplasm is known to be tightly regulated as a function of the salinity of the medium [11]. Keeping the high concentrations of ectoine necessary to balance the osmotic pressure of salt, imposes a cost on the cell, that has to mantain a level of ectoine synthesis equal to its concentration times the growth rate in order to avoid dilution. But to place this cost in the wider context of net anaplerotic fluxes, the oxaloacetate demand as a whole has to be calculated. Growth rates and intracellular ectoine concentrations, have been consistently measured for the wild type [11], and it is widely assumed that the cell composition provided by Neidhardt [1] is representative for gram-negative cells. Given this data, the demand of ectoine for growth has to be equal to its dillution rate (concentration times growth rate).The demand of OAA for aminoacid synthesis is the product of growth rate times protein content times the relative abundances of each aminoacid [33] growth at different salt concentrations can be estimated both for aminoacid and for ectoine production.

The estimate shown in figure 2 reveals that the metabolic demand for oxaloacetate in *H. elongata* is similar to that of other bacteria at low salt concentrations – e.g. 0.17 M NaCl– but the pattern quickly diverges as salt concentration increases. At the optimal interval for growth, between 0.5 M and 1 M NaCl, OAA demand reaches its peak and the cell dedicates up to 40% of the production to synthesize ectoine. As salt concentration increases even further and growth rates decline, the majority of OAA produced is destined to ectoine synthesis, thus keeping a high demand for oxaloacetate under growth rates at which other bacteria barely need any. This reflects not only a much higher demand for OAA in *Halomonas* but also a need to regulate its production differently so it can react to two different stimuli: overall growth rate and salt concentration in the medium.

**Figure 2:**
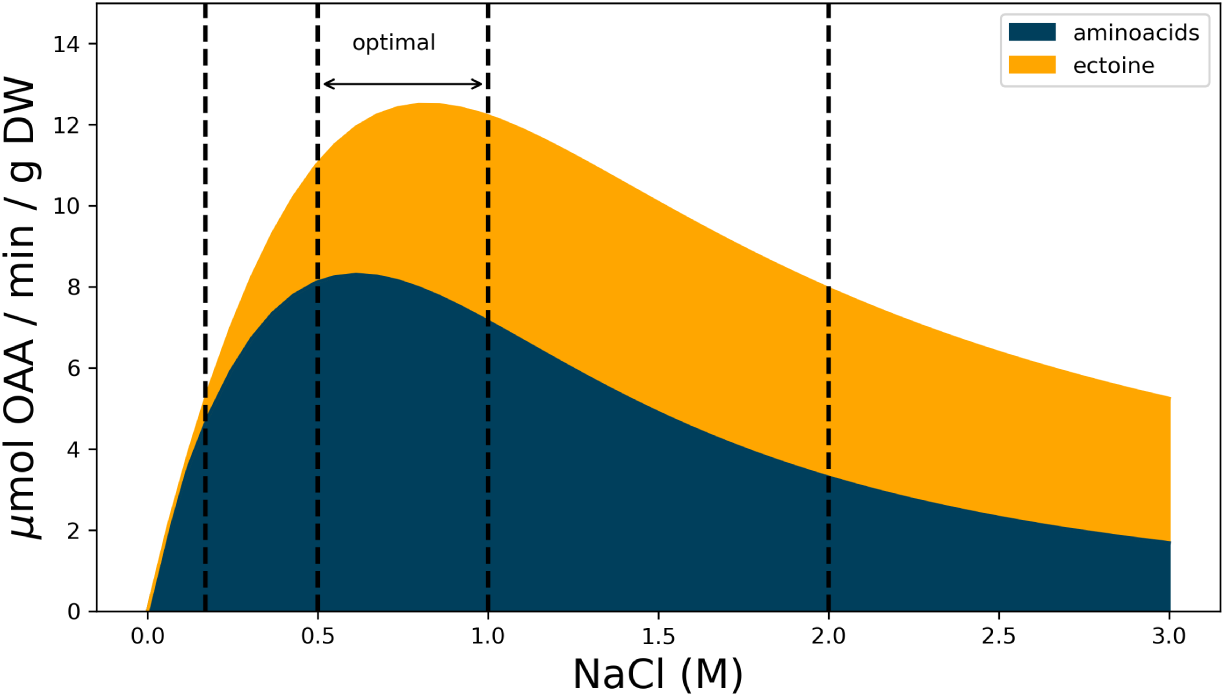
Estimated demand of OAA for ectoine and aminoacid synthesis during growth at different salt concentrations. Aminoacid demand based on protein content and aminoacid abundance similar to E. coli and ectoine demand based on reported values for *H elongata* (see text for details)

### 3.2 Only Ppc is thermodynamically favorable, Oad becomes viable at high salt

In order to meet this higher than usual demand of OAA, *H. elongata* has four enzymes that can potentially catalyze anaplerotic reactions. Only one of these four enzymes, Ppc, usually plays this role in other bacteria. The three remaining candidates, Oad, Mdh and Pck are normally considered to be thermodynamically unfavorable and thus not able to operate in the right direction within physiologically feasible concentrations of their reactants. In order to evaluate how the different environment and needs of *H. elongata* may lead to different roles for this enzymes it is important to assess how this differences have an impact on the thermodynamics of the reactions involved. As discussed above, the use of SIT enables a realistic incorporation of the sodium gradient into the thermodynamic calculations of Gibbs free energies. This is critical since for an enzyme to be able to operate *in vivo*, its free energy of reaction must be, not only negative, but high enough in absolute value that it results in an efficient use of the enzyme [22]. Standard free energies for the different reactions are often helpful to determine this, but can also be misleading, especially in this case, since standard conditions do not account for gradients across the membrane. In figure 3 we show orientative reference values for the free energies at different extracellular salt concentrations based on equation 6 and the usual rules for thermodynamic calculations of reaction-transport processes [16]. The salt dependence was calculated using physiologically realistic concentrations for intracellular sodium, phosphate and CO_2_, as well as realistic ratios for co-factors. All other substrates were assumed to be 1 mM. Of course, only the free energy of Oad changes with extracellular sodium concentration but the free energies of all potential anaplerotic enzymes are shown for comparison. Additionally a known thermodynamic bottleneck of the TCA cycle [22], malic dehydrogenase (Mdh), has been added as a reference. All free energies are calculated in the direction that leads to OAA production. The default anaplerotic reaction Ppc, shows an unsurprisingly negative free energy. Pck, which is known to be able to operate anaplerotically under extreme conditions [38] albeit very inefficiently due to unfavorable thermodynamics, provides a counterexample, with a very high, positive value. It is also noteworthy that both malic enzymes have even larger positive values, so they are very unlikely candidates for anaplerosis. Oad evolves from being very unfavorable in the absence of a sodium gradient to an enzyme able to work anaplerotically in saline environments. Interestingly, Oad is always less thermodynamically favorable than Ppc, regardless of the sodium gradient. Actually, even at high salt concentrations, Oad is still less favorable than Mdh. Thermodynamic reasons cannot therefore be invoked as an evolutionary reason to justify an anaplerotic role for this enzyme.

**Figure 3:**
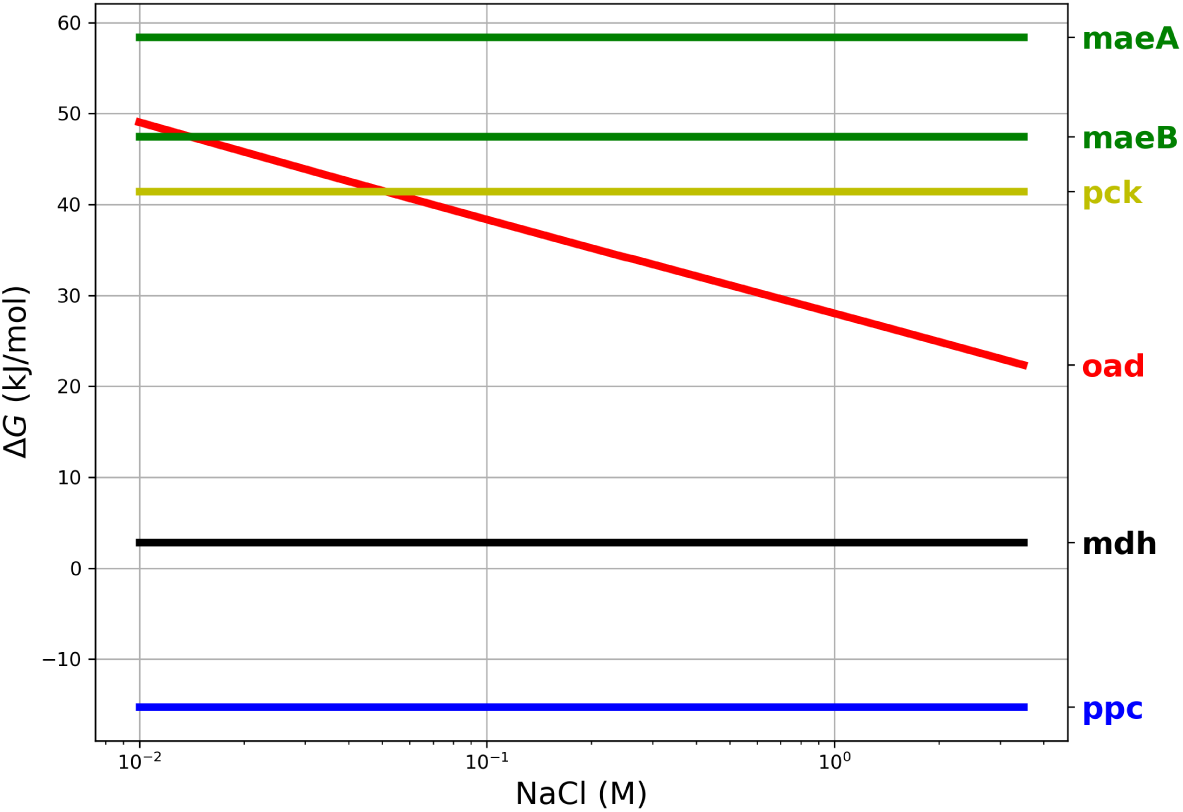
Thermodynamics of anaplerotic reactions. As a reference for *in vivo* conditions, cytoplasmic Na was fixed at 10 mM and external was varied from 0 to 3.5 M, CO _2_(total) = 10 *µ*M P = 7.5 mM cofactor ratios NADH/NAD = 0.1 NADPH/NADP = 10 ATP/ADP=10. All other metabolites were fixed at 1 mM. Malic dehydrogenase (mdh) has been added as a reference since it is known to catalyze a thermodynamically challenging reaction.

It is nevertheless important to mention that regardless of their very different thermodynamic propensities to operate in the anaplerotic direction, all enzymes are reversible within physiological concentrations of their substrates so there is no basis to discard any of them solely due to their thermodynamic properties.

### 3.3 Contributions by Pck and Mae to anaplerotic flux are severely hindered by distributed thermodynamic effects

Although all four enzymes can operate in the right direction when working in isolation, it remains to be seen whether they can carry the flux *in vivo*. In order to combine the stoichiometric and thermodynamic aspects in an integrated analysis of the relevant pathways, a TMFA model was formulated. Although focused on the TCA cycle and the anaplerotic reactions, this model must include other relevant processes such as the respiratory chain, ATPase and other processes related to sodium transport. As can be seen in figure 4, sodium plays multiple roles in this metabolic network beyond driving Oad. The electron transport chain of *H. elongata* has a sodium traslocating NADH ubiquinone oxidoreductase (Na-Nqr) [17] which couples respiration to the sodium gradient. Moreover, as many other halophilic and halotolerant bacteria, *H. elongata* has a high number of sodium-proton antiporters that can be used to built a proton gradient or to use one to pump sodium out of the cell. The different stoichiometries of the antiporters provide different degrees of coupling between proton and sodium transport just like the gears in a gearbox.

**Figure 4:**
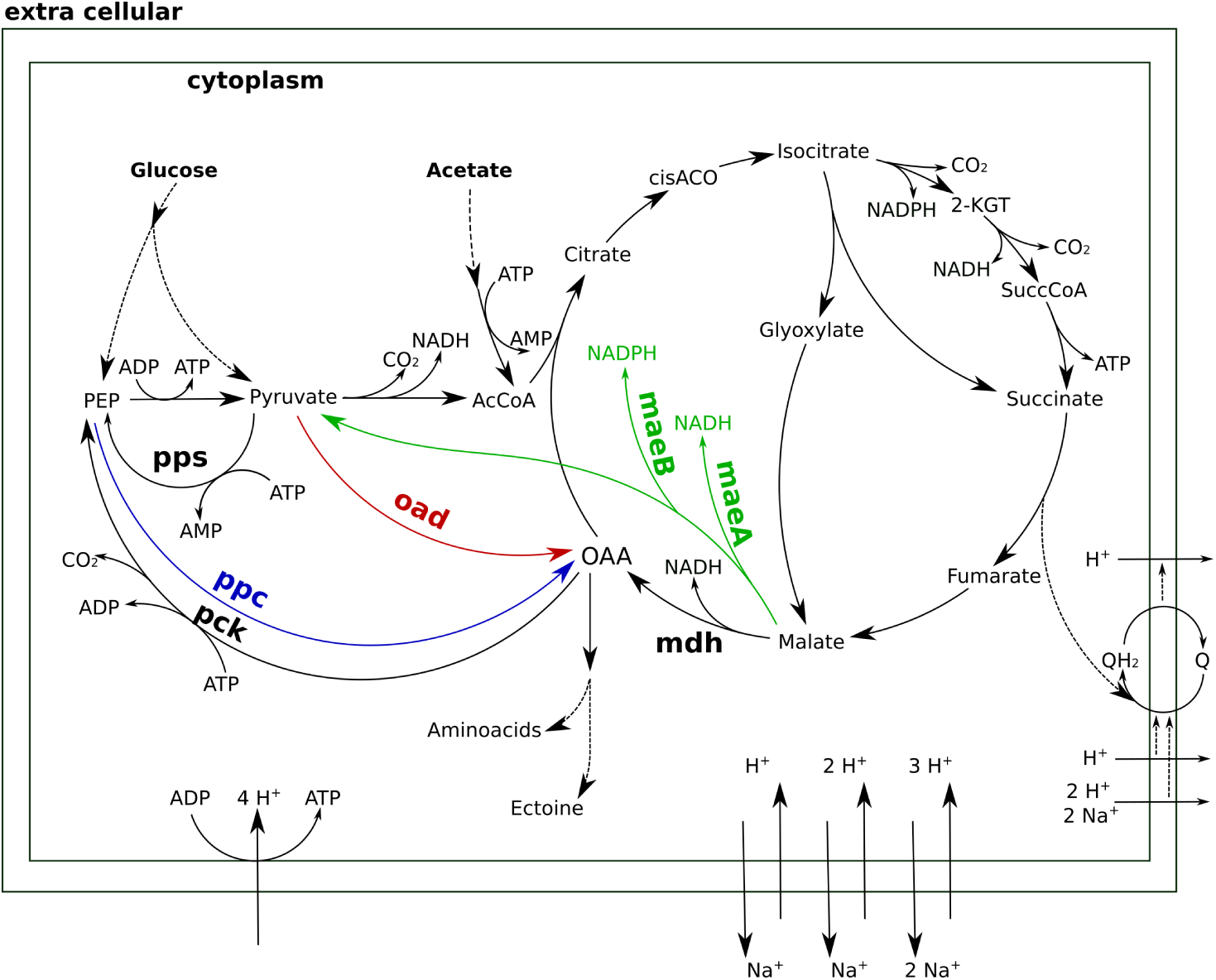
A stoichiometric model of TCA anaplerosis and related processes in *H. elongata*

Calculating the optimal flux distributions to maximize the production of Oxaloacetate points at Pck and Mae as the most advantageous anaplerotic reactions from a stoichiometric point of view, as has already been pointed out [17, 30]. In spite of their reversibility within the established *in vivo* ranges for metabolite concentrations no thermodynamically viable flux distributions could be obtained where Pck or the Malic enzymes played an anaplerotic role. In order to operate in the carboxylating direction, these enzymes require concentration ratios between their substrates and products that are incompatible with the requirements of other reactions in the network. This can be clearly seen for the malic enzyme since the low malate concentration needed to enable anaplerotic flux through Mae is incompatible with the high concentration required to drive the next enyzme in the sequence, Mdh, towards oxaloacetate production.

It is worth mentioning that computer simulations of this system often exhibit the simultaneous action of both malic enzymes in opposite directions, which results in a net transfer of electrons from NADPH to NADH. This chemically feasible solutions point to the risk faced by the cell if these two enzymes were simultaneously active. Only a strict control of the expression and activity of this enzymes can prevent them from destroying the carefully balanced redox state of the two electron currencies in *H. elongata*. Enyme assays with cell free extract have shown very low activities for Mae using NADP as cofactor [17]. No activity could be detected using NAD (personal communication, Irina Bagyan).

### 3.4 Ppc is stoichiometrically as inefficient as the glyoxilate shunt

The type of glycolysis presumed to be active in *H. elongata* is the Entner-Doudoroff (ED) pathway [18, 17]. This pathway splits glucose into pyruvate and PEP in a 1:1 proportion which poses a difficulty to channel flux through Ppc. Recovering pyruvate for anaplerosis would require its phosphorylation back to PEP, a difficult reaction that consumes two ATP equivalents and would reduce the OAA yield due to the need to divert metabolic flux for ATP production. This can be seen quantitatively by blocking all anaplerotic enzymes in the model except Ppc and then calculating the optimal flux distribution for OAA production. This simulation results in the two equivalent solutions are shown in figure 5, one makes use of Ppc for anaplerosis and the other just channels the flux through the glyoxilate shunt to achieve exactly the same yield. Thus, having Ppc as the only anaplerotic enzyme is stoichiometrically equivalent to having no anaplerotic enzymes at all.

**Figure 5:**
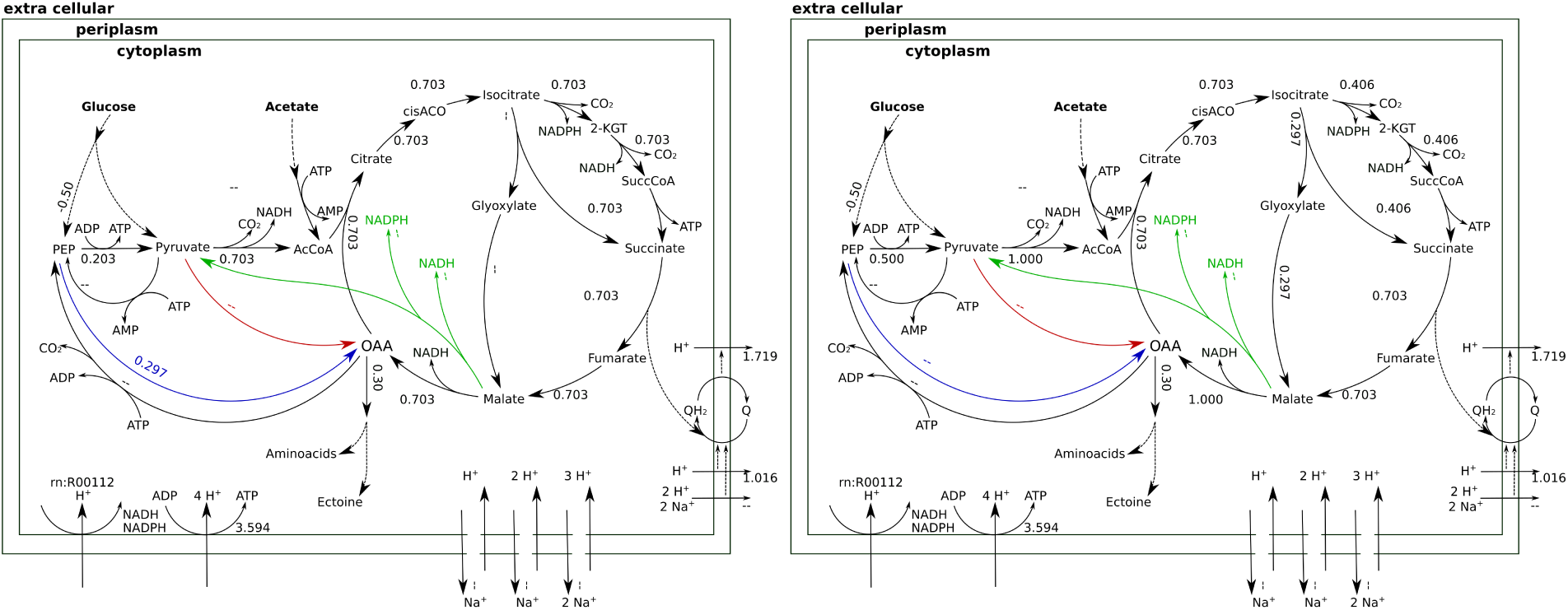
Flux distributions with optimal OAA yield indicated on the layout of figure 4. All fluxes are normalized per unit of glucose input. when Ppc is the only anaplerotic pathway available. Left: anaplerotic flux through Ppc. Right: Equivalent solution using glyoxylate cycle

When the sodium gradient allows it, Oad can carboxylate pyruvate directly to obtain higher yields of OAA. Beyond providing a stoichiometrically efficient solution, Oad ties central metabolism together with the respiratory chain and membrane transport systems into the sodium economy in the cell. The sodium influx through Oad can be a counterbalance to Na-Nqr or, when there is a net influx of sodium, be compensated with the Na/H antiporters. This adds degrees of freedom to the space of feasible flux distributions and enables the independent regulation of two parallel OAA supply fluxes.

Our analysis shows that the sodium concentration gradient across the membrane must respect a balance between two opposing forces. It must be steep enough to let Oad operate anaplerotically but not so steep that it prevents the correct operation of the electron transport chain. The tradeoff between these two selective pressures is further modulated by the viability of the different proton-sodium antiporters. Under these conditions, cytoplasmic sodium becomes a key albeit difficult to measure variable. Accurately measuring cytoplasmic sodium is an extremely difficult due to the impossibility to distinguish between its free and bound forms. Total sodium in *E. coli* has been reported to be 5 mM [32, 21] while for *H. elongata* a wide range of values has been reported between 40 and 630 mM for extracellular salt concentrations between 0.05 M and 3.4 M [36]. The range of allowed concentrations for intracellular sodium concentrations constrains the admissible values of the sodium gradient so it is a key parameter of the model. High levels of intracellular sodium will enable the operation of the electron transport chains at very high salinities and a low level can enable the operation of Oad in a low salt medium. Since it is difficult to establish this range from first principles, only experimental data can help determine this range for *H. elongata*.

### 3.5 Anaplerotic flux is distributed between Ppc and Oad with a sodium dependent ratio

In order to establish the roles of Ppc and Oad, growth experiments were done with the two mutant strains described above: *H. elongata*-PPC (Δ*ppc*::Sm) and *H. elongata*-OAD (Δ*oad*::Sm). The phenotype associated to the deletions was characterized by cultivating the wild type *H. elongata* and each of the mutants at different salt concentrations on both glucose and acetate. Growth on acetate here serves as a control in which anaplerotic flux becomes irrelevant because the incorporation of carbon into the TCA cycle occurs through the glyoxylate shunt. Therefore, any phenotype caused by an insufficiency of the anaplerotic reactions, should disappear when cells grow on acetate. A first screening was performed in a microtiter plate reader. Figure 6 shows maximum growth rates of the wild type and the different mutant strains of a representative experiment. The growth rates are comparable between all strains in most conditions. The variability between strains is normally within the inter experiment variability of the wild type. Only the *H. elongata*-OAD strain shows a significant decrease in maximum growth rate at 1 M salt compared to the wild type. Repetition of the experiment under this condition with a higher number of replicates did not reproduce this difference in growth.

**Figure 6:**
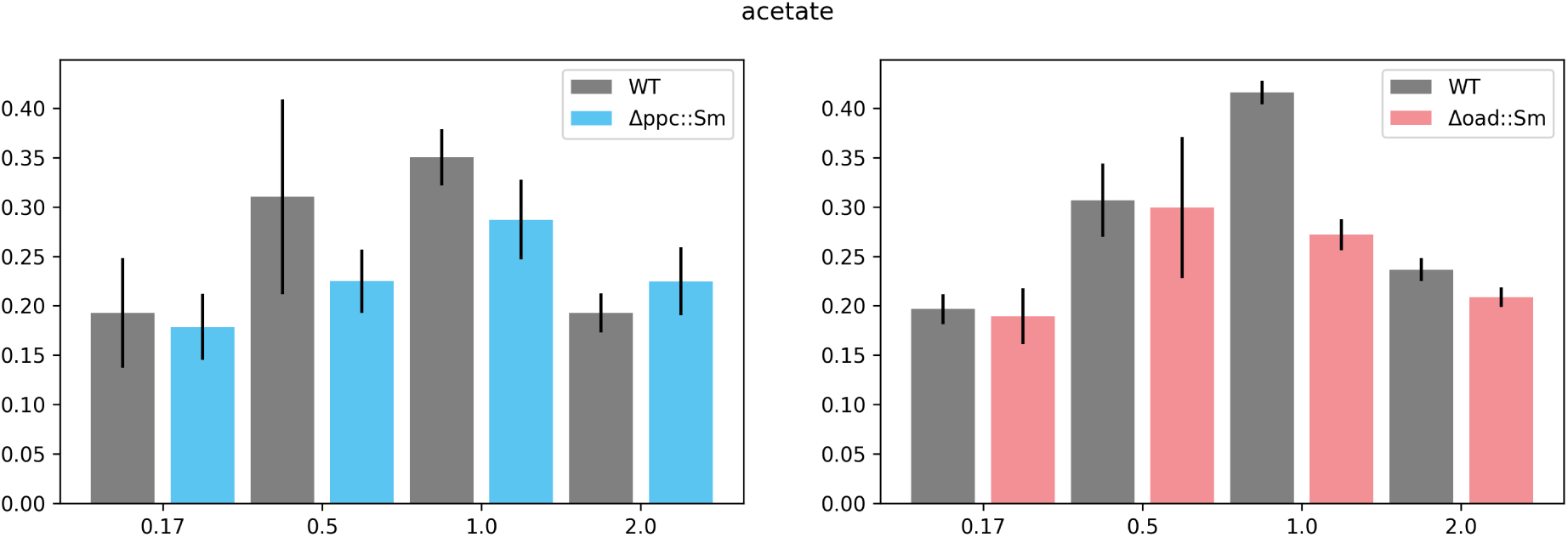
Growth of *H. elongata* on Acetate. Cells of each of the mutant *H. elongata* strains were grown at 30 *°* C in mineral salt medium (MM63) with NaCl (salt) at a concentration of 0.17 M, 0.5 M, 1 M, and 2 M, respectively, and with acetate as sole carbon source. Each of the experiments compared maximum growth rates of a mutant strain (*H. elongata*-PPC in blue and *H. elongata*-OAD in red) in comparison to the wild type (WT black).

The equivalent experiment with glucose as carbon source is shown in figure 7 and exhibits similar degrees of variability with all strains reaching similar growth rates in all cases except two. The strain lacking Ppc exhibits a severe inhibition of growth at low salt concentrations (0.17 M) whereas the strain lacking Oad shows similar difficulties at high salt concentration (2 M).

**Figure 7:**
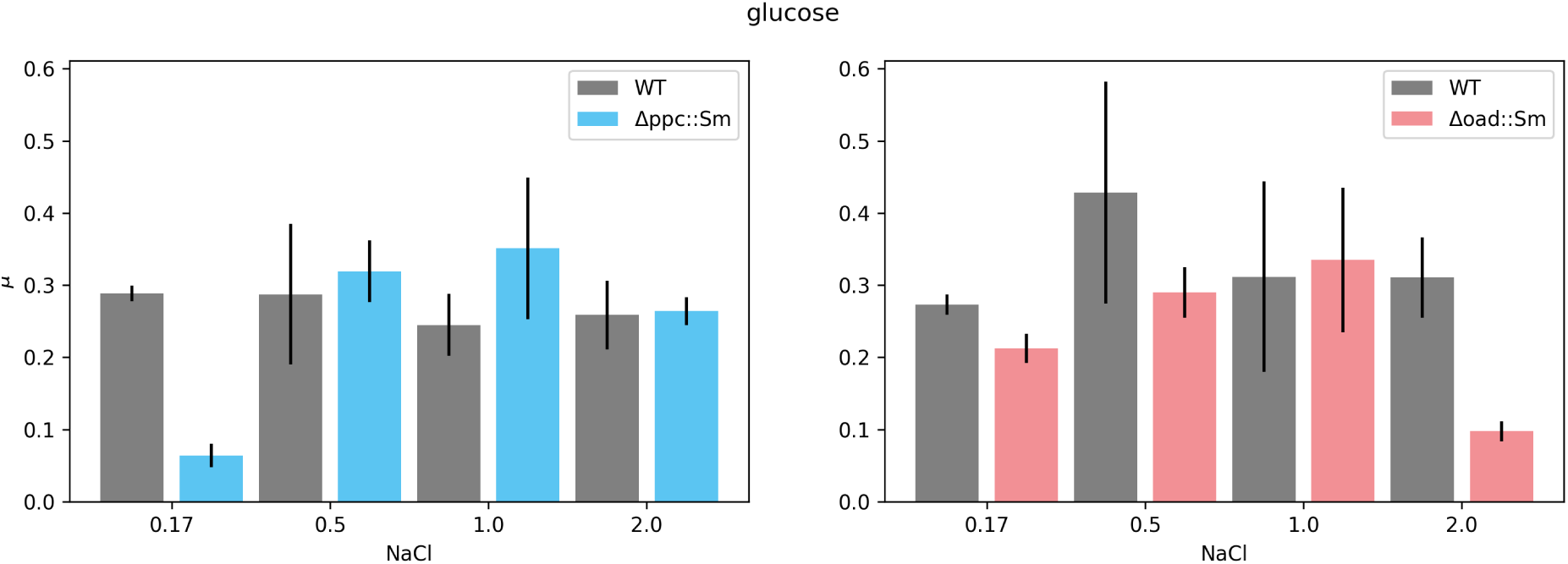
Growth of *H. elongata* on Glucose. Cells of each of the mutant *H. elongata* strains were grown at 30 *°* C in mineral salt medium (MM63) with NaCl (salt) at a concentration of 0.17 M, 0.5 M, 1 M, and 2 M, respectively, and with glucose as sole carbon source. Each of the experiments compares maximum growth rates of a mutant strain (*H. elongata*-PPC in blue and *H. elongata*-OAD in red) in comparison to the wild type (WT black).

A further round of experiments was carried out in flasks for confirmation of the most significant phenotypes observed during the micro-plate screening. In most cases, the flask experiments yielded a more accurate picture of what was already indicated by the previous screening. *H. elongata*PPC growing in a medium with 0.17 M salt exhibited similar growth rates as the wild type when growing on acetate (0.15 ± 0.02 vs 0.18 ± 0.002) but experienced difficulties when growing on glucose (0.07 ± 0.04 vs 0.27 ± 0.008). Comparison of growth rates on glucose at high salt, however, provide a finer comparison between the three strains. Besides the dramatic drop from a growth rate of 0.25 ± 0.01 *h*^−1^ in the wild type to 0.08 ± 0.003 *h*^−1^ in *H. elongata*-OAD, a smaller decrease to a growth rate of 0.15 ± 0.01 *h*^−1^ is shown in the Ppc lacking strain, which is only 0.6 times as fast as the wildtype. Again, increased lag phases for both mutant strains on glucose were observed especially during growth in their respective inhibitory salt condition. Furthermore, the deletion of ppc seems to generate a rather unstable mutant showing a higher variance in overall growth behaviour.

These phenotypes shed some light on the lower limit for intracellular sodium concentration. Thermodynamic simulations indicate that intracellular sodium levels of 10 mM or lower would enable anaplerosis through Oad at 0.17M so this concentration is either unreachable for the mutant strain *H. elongata*-OAD or has a deleterous effect on the cell.

### 3.6 Robustness of ectoine levels at high salt

The ectoine content of cells growing at 2 M NaCl was measured in wild type *H. elongata* as well as in the various mutants obtained in this study and is shown in figure 8. Low salt conditions were not included in this analysis because the levels of ectoine in the cell under such conditions are too low to be measured reliably. For these experiments, oxaloacetate demands – and therefore total anaplerotic flux – can readily be calculated. The different strains showed very similar ectoine concentrations, even when growing on different carbon sources, which is consistent with the strict regulation expected. The largest deviation from the wild type was the *H. elongata*-OAD strain, which with a roughly 25% lower ectoine content only grows 50% as fast. The estimate of OAA demands shows how the strain without Oad has difficulties keeping up the anaplerotic flux, being able to reach only 51% of the anaplerotic flux of the wild type. It is noteworthy that the strain without Ppc cannot reach the full flux either, totaling 75% of that of the wild type. These results point to a shared anaplerotic role between Oad and Ppc even at fairly high salt concentrations.

**Figure 8:**
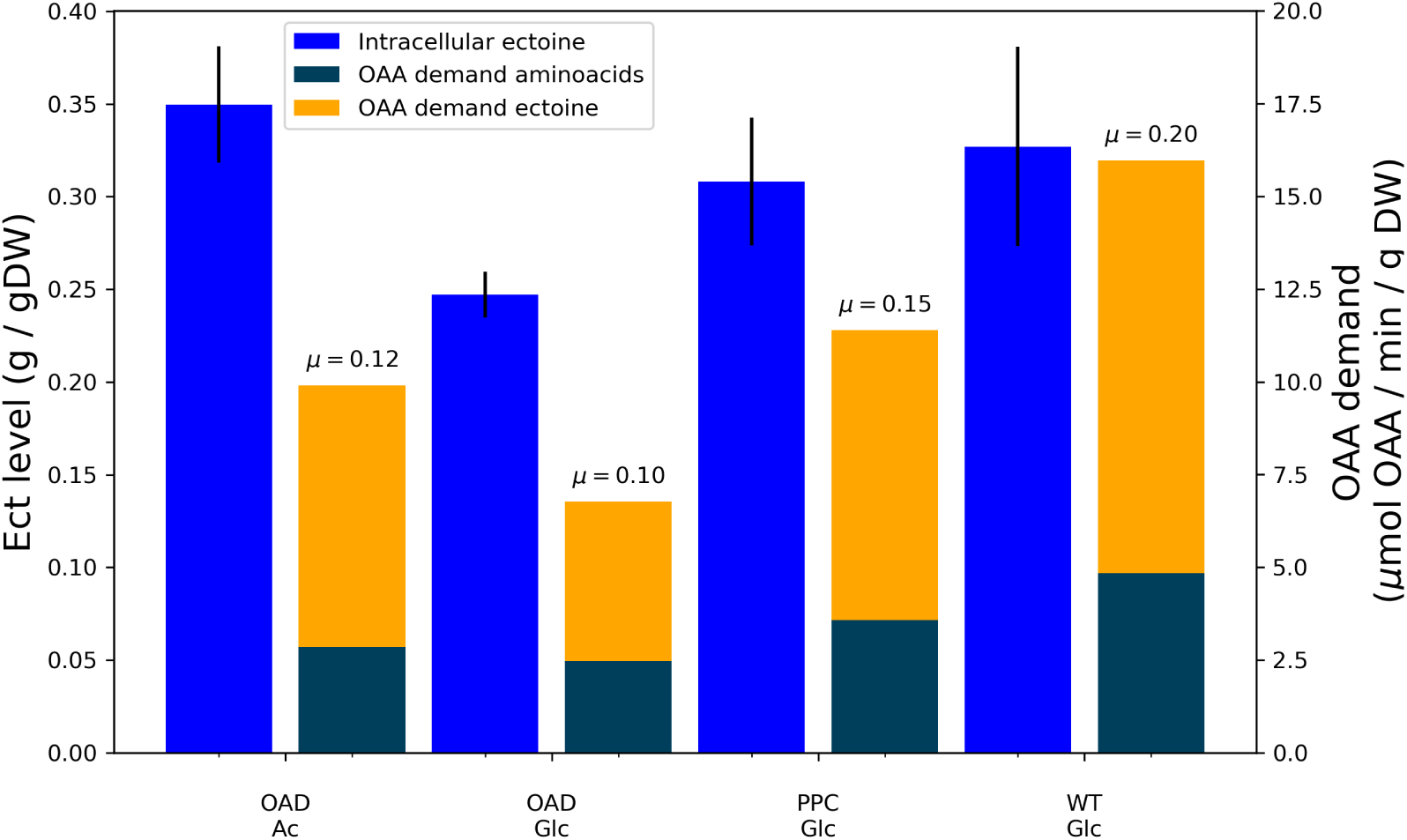
Ectoine content (left bars) and oxaloacetate demand for aminoacid and ectoine synthesis (right bars stacked) in different strains at 2M salt. Growth rates indicated on top of oxaloacetate demand bars

## 4 Discussion

The results of this study clearly indicate that Oad can operate as an anaplerotic enzyme as was hypothesized by Kindzierski [17]. Furthermore, it can even carry most of the necessary flux for normal growth and ectoine production in the absence of Ppc, as long as the salt concentration is high enough. Part of the flux, however, seems to go through Ppc even at salt concentrations where Oad by itself would be able to channel the totality of the anaplerotic flux.

The co-existence of alternative anaplerotic enzymes, and of parallel pathways in general, is frequent among bacteria [29]. This can be understood in evolutionary terms as a source of robustness through redundancy, a way to ensure flexibility by providing alternatives that work well under different conditions. An example of this are the high and low affinity pathways for ammonia assimilation, which guarantee an efficient incorporation of nitrogen into amino acids regardless of the concentration of ammonia [34, 31]. But metabolic alternatives can also be a regulatory necessity as is the case with some isoenzymes. The parallel operation of several enzymes at the beginning of a branched pathway – e.g. amino acid synthesis – enables the independent inhibition of each by a different end-product. The unusual architecture of the anaplerotic reactions in *H. elongata* seems to obey to a combination of all of the above. Ppc and Oad offer alternative pathways to guarantee an anaplerotic flux under different conditions. That these conditions overlap is shown by the fact that a mutant lacking either enzyme can achieve the same growth rate as the wild type at some salt concentrations. Moreover, as this work shows, low salt concentrations render Oad as an inviable option, so the presence of Ppc widens the range of salinities in which *H. elongata* can thrive. Finally, the anaplerotic reactions are a critical part of two different metabolic functions providing precursors for both growth and salt tolerance. Since the environmental factors that condition these two functions are independent from one another, the coexistence of different enzymes enables a more flexible regulatory scheme where different enzymes can be modulated by different factors. The sodium dependence of Oad guarantees an increase in anaplerotic flux to support increased ectoine production at higher salt while Ppc can respond to an increased demand of OAA for aminoacid biosynthesis. Further investigations on this topic will surely provide valuable insights and improve our understanding of the design principles that shape this part of bacterial metabolism.

## 5 Acknowledgements

This work has been funded by the German Federal Ministry of Education and Research (BMBF) through projects OpHeLIA (0316197) and HOBBIT (031B03).

